# *rustybam:* a composable toolkit for alignment analysis and visualization with *SafFire*

**DOI:** 10.64898/2026.02.16.706142

**Authors:** Mitchell R. Vollger

## Abstract

**Summary:** We present *rustybam*, a Rust-based command-line toolkit for CIGAR-aware manipulation of PAF and BAM alignments, and *SafFire*, a browser-based tool for interactive genome comparison visualization with annotation overlays. *rustybam’s* composable subcommands provide overlap resolution, coordinate liftover, alignment splitting at structural variants, scaffolding, and identity statistics.

**Availability and implementation:** Both tools are open source under the MIT license. *rustybam* is available via Bioconda (https://anaconda.org/bioconda/rustybam), crates.io, and GitHub (https://github.com/vollgerlab/rustybam). *SafFire* is hosted at https://vollgerlab.com/SafFire/ with source code at https://github.com/vollgerlab/SafFire.

## Introduction

Telomere-to-telomere (T2T) assemblies of diploid human genomes and pangenomes have made whole-genome pairwise alignment a routine step in comparative genomics (Nurk et al. 2022; Liao et al. 2023). The minimap2 aligner (Li 2018) established the Pairwise mApping Format (PAF), now a de facto standard for representing pairwise alignments. However, whole genome alignment presents challenges at copy number and structurally variable loci where supplemental alignments are required. In these regions, aligners (including minimap2) can produce overlapping alignments in which the same query bases are mapped to multiple target positions across different supplemental alignments. Without resolution, these overlaps inflate coverage estimates, confound breakpoint identification, produce misleading visualizations, and confound liftover operations.

A rich ecosystem of tools supports alignment post-processing. paftools.js provides alignment, simple variant calling, statistics, and liftover operations, and wgatools (Wei et al. 2025) offers format conversion, visualization, filtering, and more. Additional options are available for visualization including D-GENIES (Cabanettes and Klopp 2018), mummerplot (Marçais et al. 2018), ModDotPlot (Sweeten, Schatz, and Phillippy 2024), StainedGlass (Vollger, Kerpedjiev, et al. 2022), plotsr (Goel and Schneeberger 2022), and SVbyEye (Porubsky et al. 2025).

We present *rustybam*, a Rust-based toolkit of composable subcommands for CIGAR-aware manipulation of PAF and BAM alignments, and *SafFire*, a browser-based visualization tool for interactive genome comparisons with annotation overlays. *rustybam’s* subcommands chain through Unix pipes, allowing users to build tailored workflows. While *rustybam’s* functionality overlaps with many existing toolkits, it provides two unique features: *rustybam liftover* performs coordinate conversion while maintaining alignment information, and *rustybam trim-paf* uses dynamic programming to prevent query sequence from aligning to more than one position in the target genome. *SafFire* renders the resulting alignments as interactive miropeats-style (Parsons 1995) ribbon plots with annotation overlays and URL-based view sharing. Both tools have been used extensively in T2T Consortium (Nurk et al. 2022; Vollger, Guitart, et al. 2022; Rautiainen et al. 2023; Rhie et al. 2023) and Human Pangenome Reference Consortium (Liao et al. 2023; Vollger et al. 2023) publications.

## Implementation

### rustybam

*rustybam* is written in Rust and distributed via Bioconda, crates.io, and pre-compiled binaries. Its core design principle is that all coordinate operations preserve CIGAR string integrity: when an alignment is trimmed, split, or lifted over, the CIGAR string is updated to reflect the exact change, ensuring that downstream identity calculations and variant calls remain accurate. rustybam accepts both cg (CIGAR) and cs (difference string) tags, preserving individual base substitutions and indel sequences that cannot be reconstructed from CIGAR alone.

The toolkit provides subcommands invoked as *rb <subcommand>* (or equivalently *rustybam <subcommand>*), designed to chain through Unix pipes. The key subcommands include:

***liftover*** projects target-coordinate BED intervals onto query coordinates (or vice versa with *--qbed*) through PAF alignments. Unlike other liftover tools (paftools.js, UCSC liftOver, etc.), which return lifted coordinates only, *rb liftover* outputs a trimmed PAF record with an updated CIGAR string spanning exactly the lifted region. This design enables direct composition with other *rustybam* subcommands: for example, piping *rb liftover* into *rb stats* computes per-region alignment identity in a single command, allowing users to assess sequence conservation across lifted intervals without additional tools.

***trim-paf*** resolves overlapping query alignments that arise at duplications and inversion boundaries. The algorithm loads all PAF records, identifies query-coordinate overlaps, and uses cumulative-score maximization with configurable match, mismatch, and indel scores to find the optimal split point within each overlapping CIGAR string. The process iterates from the largest overlap until no query base is aligned more than once, producing clean, non-overlapping alignments suitable for breakpoint analysis and visualization (Fig. 1). Because the trimming operates directly on the CIGAR string, the resulting records retain accurate alignment coordinates and identity statistics.

**Figure 1.**
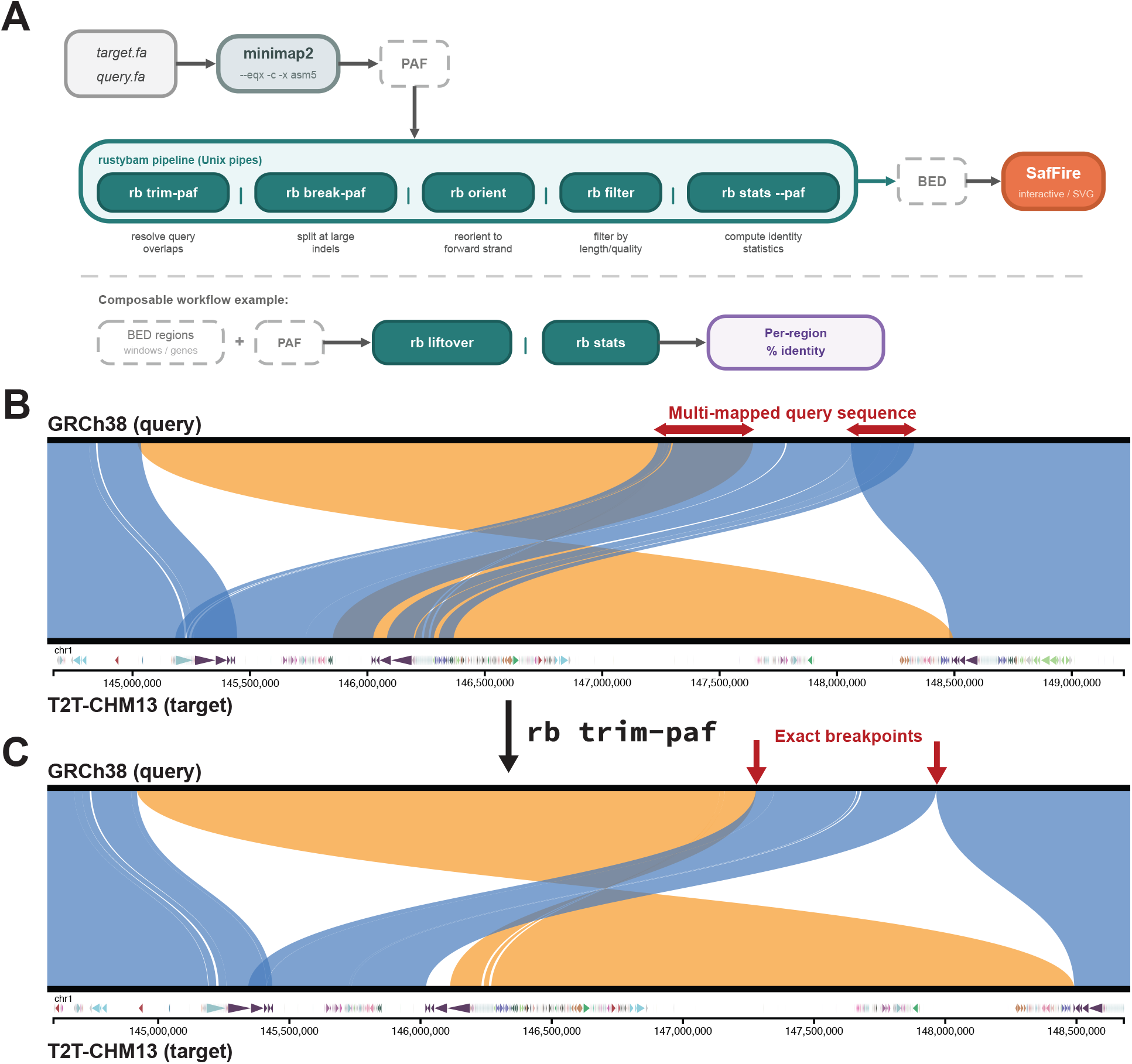
The *rustybam*–*SafFire* pipeline. **(A)** Schematic of the data processing pipeline from minimap2 PAF through *rustybam* subcommands to *SafFire* visualization. **(B)** *SafFire* visualization of the NOTCH2NL locus on chromosome 1, comparing T2T-CHM13v2.0 (target) and GRCh38 (query). Blue ribbons indicate forward alignments; orange ribbons indicate inversions. BED annotations show segmental duplications annotated by DupMasker. **(C)** The same locus after applying *rb* t*rim-paf*, showing resolution of overlapping alignments at duplication boundaries visible in (B).

***break-paf*** splits alignment records at insertions or deletions exceeding a user-defined threshold (e.g., 5,000 bp). This transforms large alignments into fine-grained alignment segments for downstream visualization.

***orient*** reorients query contigs so that the majority of aligned bases are in the forward orientation, simplifying visualization and downstream analysis. An optional scaffold mode merges multiple query contigs per target into a single pseudo-scaffold ordered by target position.

***stats*** computes per-alignment identity and coverage statistics directly from CIGAR strings into BED format with identity annotations, providing the input format used by SafFire.

An additional 11 subcommands provide utility operations including PAF filtering, coordinate inversion, sequence extraction, nucleotide frequency calculation, format conversion, and assembly statistics.

A key design goal is that subcommands compose through Unix pipes to build analysis workflows from simple operations. Because each subcommand reads and writes PAF (or BED) through standard streams, users can construct pipelines tailored to specific analyses. For example, lifting gene annotations through an alignment and computing per-gene identity requires only:

rb liftover --bed genes.bed aln.paf | rb stats --paf

Similarly, you can combine liftover with overlapping windows to create sliding measures of percent identity, e.g.:

rb liftover --bed <(bedtools makewindows -g ref.chrom.sizes -w 10000 -s 1000) aln.paf | rb stats --paf

### SafFire

*SafFire* is a browser-based visualization tool that renders miropeats-style (Parsons 1995) genome comparison views as an interactive graphic. It is implemented entirely in client-side JavaScript using D3.js (Bostock, Ogievetsky, and Heer 2011), requiring no software installation—users simply visit the hosted instance or serve the files locally with any static HTTP server. *SafFire* complements tools like SVbyEye (Porubsky et al. 2025) and plotsr (Goel and Schneeberger 2022), which provides miropeats-style plots within R/python workflows, by offering a browser-native experience with interactivity and URL-based view sharing.

*SafFire* takes as input the BED-format output of rb stats --paf, typically after processing through the *rustybam* pipeline:

minimap2 --eqx -c -x asm5 ref.fa query.fa \

| rb trim-paf | rb break-paf -m 5000 \

| rb orient | rb filter --paired-len 100000 \

| rb stats --paf > input.bed

Each alignment is rendered as a colored ribbon connecting target and query coordinates: blue for forward alignments, orange for reverse (inversions). Ribbon opacity encodes percent identity, and a percent-identity track below the target axis provides an alignment quality profile. Users can zoom and pan along the target axis, select individual target and query contigs, and click positions to copy genomic coordinates.

Key features include: (i) BED annotation overlays for both target and query sequences, enabling display of gene annotations, centromere satellite classifications, segmental duplication colors, or any custom annotation; (ii) UCSC Genome Browser integration that synchronizes a browser snapshot to the current viewport; (iii) SVG export for publication-quality figures; and (iv) URL-based state sharing via hash parameters (\#ref=&query=&pos=), allowing specific views to be bookmarked and shared. *SafFire* includes pre-loaded datasets with alignments to CHM13 and GRCh38 references, including HPRC and non-human primate assemblies.

## Results

### Performance

To evaluate *rustybam’s* performance (v0.2), we benchmarked key subcommands on whole-genome PAF alignments generated by minimap2 (--eqx -c -x asm5) between CHM13v2 and GRCh38 (1,460 records, 69 MB). For less intensive subcommands, we used a 20×-concatenated version of this PAF (29,200 records, 1,372 MB) to ensure wall times exceeded measurement noise. All benchmarks used hyperfine with 2 warmup and 5 timed replicates on a MacBook Pro (Apple M4, 64 GB RAM) running single-threaded (-t 1).

*trim-paf* resolved all overlapping alignments across the whole genome in 8.9 s (±0.1 s) on the 1× input. On the 20× input, break-paf completed in 8.6 s (±0.4 s) at a 1,000 bp threshold and 10.5 s (±1.1 s) at 5,000 bp, orient in 8.6 s (±0.5 s), and filter in 8.6 s (±0.5 s). For alignment statistics, rb stats and paftools.js stat showed comparable performance: rb stats was slightly faster on the 1× input (155 ms vs.\ 207 ms for paftools.js) while paftools.js was slightly faster on the 20× input (2.7 s vs.\ 3.4 s for rb stats), likely due to their different parsing approaches.

### Liftover accuracy

We compared *rb liftover* against paftools.js liftover on 14,565 BED regions lifted through an alignment of T2T-CHM13 to GRCh38 (using paftools.js -l 0 -q 0). Both tools successfully lifted the same 14,274 regions. For the 14,733 coordinate pairs output by paftools.js, 14,666 (99.5%) had identical coordinates and the remaining 67 (0.5%) differed by exactly 1 bp at the end coordinate, where paftools.js places the end coordinate 1 bp into an insertion. paftools.js was faster than *rustybam* liftover due to their different output strategies: paftools.js performs a block-based coordinate lookup and returns lifted coordinates, while rb liftover walks and trims the full CIGAR string to produce a truncated alignment record that can be piped directly into downstream PAF analysis such as *rb stats*.

### Biological application

Figure 1 demonstrates the *rustybam*–*SafFire* pipeline applied to the *NOTCH2NL* locus on chromosome 1, a region of medical importance containing segmental duplications that differ between GRCh38 and CHM13 (Mefford et al. 2008; Vollger, Guitart, et al. 2022). The *trim-paf* step resolves overlapping alignments at duplication boundaries, preventing double-counting in identity estimates and identifying base-pair precise breakpoints. The *SafFire* visualization reveals the complex pattern of repeated and inverted duplications, with BED overlays highlighting segmental duplication classifications.

## Conclusion

*rustybam* and *SafFire* extend the comparative genomics toolkit with CIGAR-aware alignment manipulation and interactive visualization. *rustybam’s* core contributions—overlap resolution (*trim-paf*) and alignment-preserving liftover (*liftover*) complement existing tools such as paftools.js and wgatools. *SafFire* adds interactivity with annotation overlays and URL-based view sharing alongside visualization tools like D-GENIES, SVbyEye, and ModDotPlot, and plotsr. Both tools have been used in T2T Consortium and HPRC publications (Nurk et al. 2022; Vollger, Guitart, et al. 2022; Rautiainen et al. 2023; Rhie et al. 2023; Liao et al. 2023; Vollger et al. 2023), accumulating over 80,000 downloads on Bioconda and crates.io.

## Acknowledgements

The authors thank A. Lo for help in editing this manuscript and members of the T2T Consortium and Human Pangenome Reference Consortium for feedback and testing.

## Data availability

All software is open source (MIT license). *rustybam* is archived on Zenodo (DOI: 10.5281/zenodo.4654875) and *SafFire* on Zenodo (DOI: 10.5281/zenodo.5765964). Example datasets and a hosted *SafFire* instance are available at https://vollgerlab.com/SafFire/.

